# APCalign: an R package workflow and app for aligning and updating flora names to the Australian Plant Census

**DOI:** 10.1101/2024.02.02.578715

**Authors:** Elizabeth Wenk, William Cornwell, Anne Fuchs, Fonti Kar, Anna Monro, Hervé Sauquet, Ruby Stephens, Daniel S Falster

**Affiliations:** Evolution & Ecology Research Centre, The University of New South Wales Sydney, NSW, Australia; Centre for Australian National Biodiversity Research (a joint venture between Parks Australia and CSIRO), Canberra, ACT, Australia; National Herbarium of NSW, Botanic Gardens of Sydney, Mount Annan, NSW, Australia; School of Natural Sciences, Macquarie University, Ryde, NSW, Australia

## Abstract

Here we present “APCalign”, an R package and accompanying browser-sourced application to align and update scientific names for Australian vascular plants to the most likely currently accepted name using the Australian Plant Census (APC) or a name in the Australian Plant Names Index (APNI). Scientific names are the label assigned to unique taxon concepts by the scientific community, but this common terminology is most useful if a taxon concept is consistently referred to by the same name. These links can be broken due to either spelling mistakes or taxonomic changes. Automated tools are required to resolve taxon lists, aligning and updating long lists of possibly erroneous scientific names to the most likely currently accepted names. It is essential that tools specific to the APC/APNI be developed, as these lists specify an endorsed national-level nomenclature used in government legislation and include the uniquely Australian concept of phrase names, absent in global taxonomic datasets. To align input names to names within the APC or APNI, “APCalign” works progressively through a sequence of checks that combine different permutations of the input name, exact versus fuzzy matches, matches that consider the entire name input versus a subset of words, and character strings that indicate a name can only be resolved to a genus or family. The aligned names are then, when possible, updated to a currently accepted taxon concept within the APC. This package should facilitate all research outputs that require diverse scientific name lists to be merged or outdated lists to be updated.

## Introduction

Taxonomic names and lists are essential in modern e-research infrastructure, enabling us to link concepts about what taxa exist to information about where they occur, how they are related to one another, and what traits they have. These linkages are easily broken or missed, however, due to inconsistencies in the delineation and naming of taxon concepts among different datasets (Franz and Peet 2009; Sandall et al. 2023). The scientific name applied at a given point in time indicates a snapshot of our understanding of evolutionary relationships within a taxonomic group. However, names change, as understanding of what morphological or genetic features circumscribe a specific taxon concept are refined. Scientific names are also easily misspelt or mistyped. When dealing with large numbers of taxa, such errors are hard to spot. To maintain the unique link between a scientific name, the associated taxon concept, and associated data, diverse users must be able to automatically align and update a list of possibly inaccurate or outdated scientific names to the currently accepted scientific name.

A range of software tools have been developed to resolve taxonomic mismatches (to the extent possible), each referencing one or more taxonomic datasets (Grenié et al. 2023; Schellenberger Costa et al. 2023), but as yet, none are specific to the current Australian National Species Lists (auNSL) for vascular plants (https://biodiversity.org.au/nsl/). Typically, these tools input a list of species names, returning a list of said names matched to the most likely currently accepted name. Working out the best or most likely match for a species name can be a computationally intensive task; still, modern tools have found several approaches to do this with reasonable speed and measurable accuracy. Different tools also have different features (Figure 1) and are presented in a range of formats, from more accessible browser-based tools, such as the Taxonomic Name Resolution Service (TNRS; Boyle et al. 2013) to more specialised R packages, such as the “taxize” R-package (Chamberlain et al. 2020; Chamberlain and Szöcs 2013). While all existing global tools do handle names of Australian taxa, the taxon lists and taxonomy in these global resources can differ from our National Species Lists, as they are managed by different authorities, each with their own missions and guidance councils (Garnett et al. 2020; Schellenberger Costa et al. 2023).

**Figure 1.**
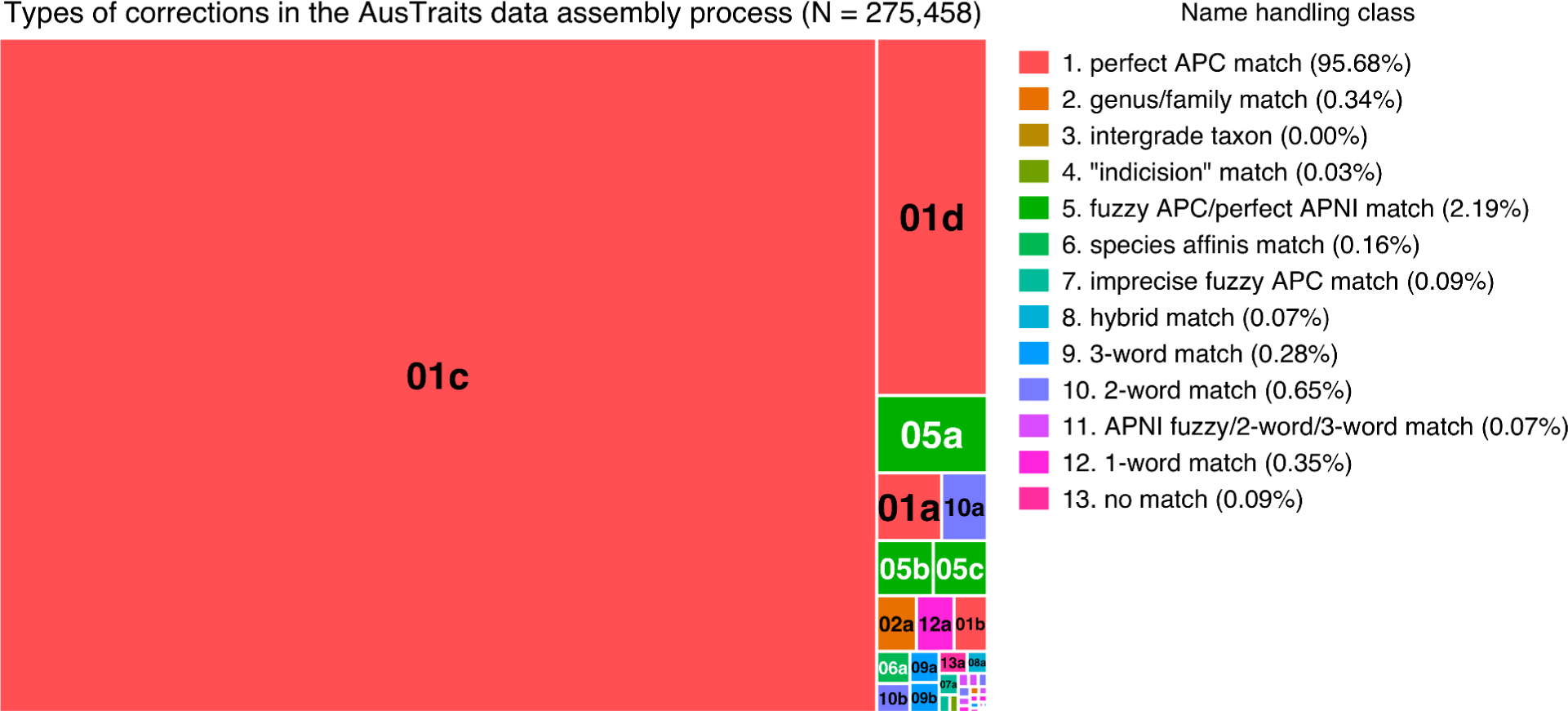
Frequency of different name handling classes found during alignment of 45,337 names from 362 raw inputs to the AusTraits compilation (Falster *et al.* 2021). See Table 4 for details of each class and subclass used for handling names.

**Figure 2.**
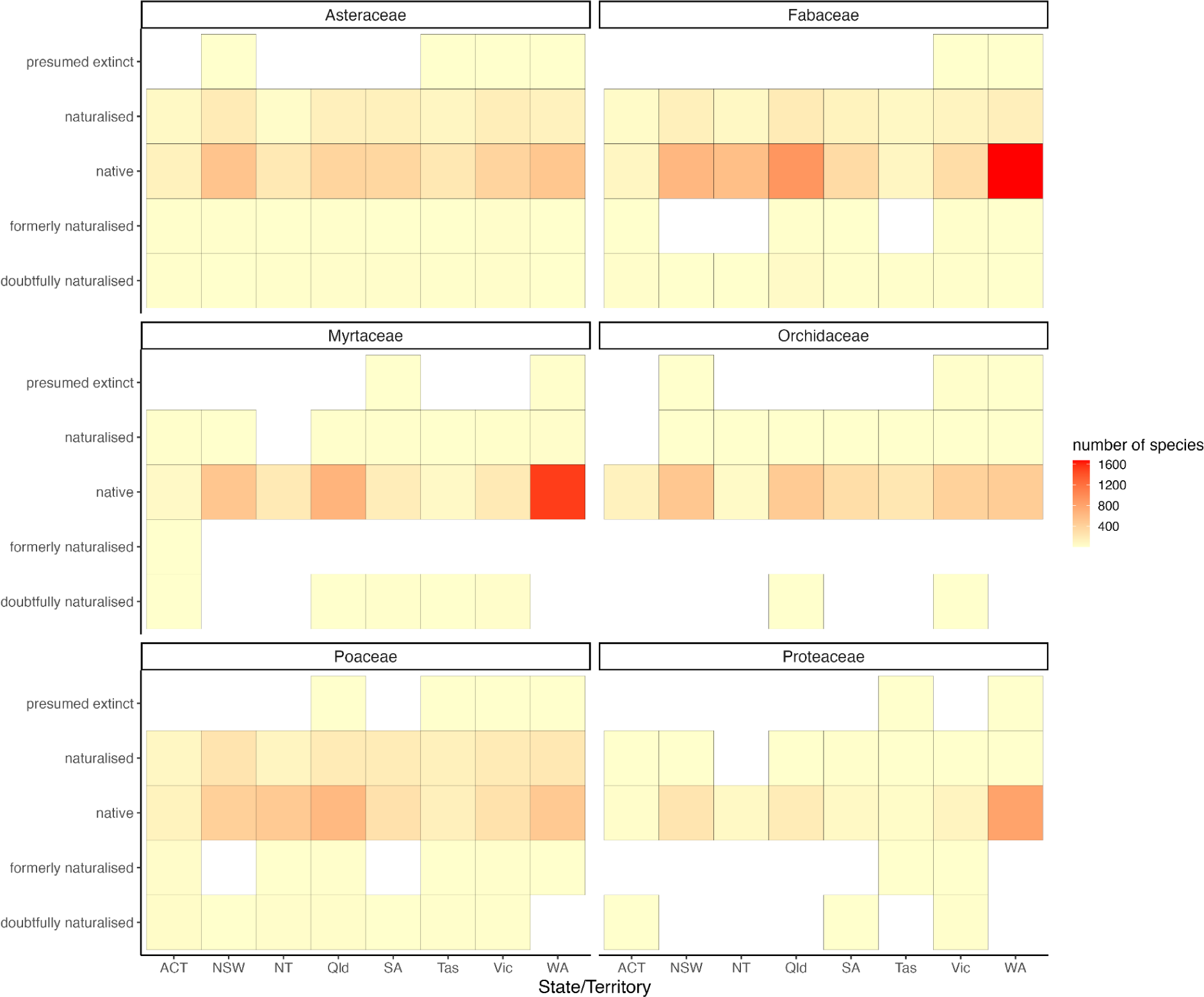
Distribution of the number of vascular plant species in several major Australian families, split by family, state/territory, and establishment means. Note that individual species may appear in different states with different statuses, and the striking variation in geographic diversity patterns across families.

In Australia, the national taxonomic standard for vascular plants is the Australian Plant Census (APC), supported by a comprehensive list of plant names documented in references: the Australian Plant Names Index (APNI). The APC is endorsed by the Council of Heads of Australasian Herbaria (CHAH), with taxonomy agreed upon by consensus. The APC is updated on a regular basis, following a review of both newly described plant taxa and taxonomic revisions, first by a working group comprising representatives from all major Australasian herbaria, followed by ratification by CHAH. The working group reviews taxon concepts extracted from recent taxonomic publications, ensuring Australia’s botanical community considers newly published names and changed taxon circumscriptions robust. The names documented in APNI and accepted by APC may be out-of-step with those used in international checklists, as each checklist may deem different scientific names or taxonomies as “current”. Having a national checklist is also essential for Australia due to the inclusion of large numbers of phrase names within the APC/APNI that are excluded from international checklists.

Phrase names are assigned to unique taxa that are yet to be formally described, a nomenclatural process that is particular to Australian plant taxonomy (Barker 2005). These names follow a specific convention: the generic name, a rank indicator, a geographic or morphological descriptor, the collector’s name and a number representing a herbarium voucher for the new taxon concept. Moreover, as our official national list, the APC is the natural (sometimes mandated) focus for documenting and linking data for Australian vascular plants.

Here we provide a new software tool, “APCalign”, that can quickly match plant names to accepted taxon concepts in the APC and to names in APNI. “APCalign” can function both as an R package for scientific and technical users, and in a browser interface for lay users. “APCalign” offers a two-step process for creating a list of names matched to the APC/APNI. In the first step, algorithms align each name input to the best match of any name within the APC/APNI, allowing matches to the infraspecies, species, genus, or family level, as is appropriate. In the second step, the aligned names that have been matched to a scientific name within the APC are updated to a currently accepted taxon concept within the APC. Our implementation builds on best practices for global taxonomic resources, as well as additional enhancements (Table 1). It includes a well-considered and well-tested sequence of algorithm matching, including both complex string matches and fuzzy-matching, to maximise meaningful links. The taxonomic updating functions handle taxonomic uncertainties. For scientific names linked to taxon concepts accepted by APC, the final table adds distribution and native status. In total, ten functions are exported by “APCalign” (Table 2), including functions to download taxonomic resources, functions to clean, align and update names to a name recorded in the APC/APNI, and functions to tabulate information about taxon distribution, diversity, and native status. By working directly with Australia’s National Species Lists for vascular plants, “APCalign” fills an important gap in our biodiversity e-resources.

**Table 1.** Comparison of features in “APCalign” and existing name matching tools.

|  | APCalign | taxize | TNRS | kewr |
| --- | --- | --- | --- | --- |
| Batch processing | X | X | X | X |
| Fuzzy matching | X | X | X |  |
| Adjustable fuzzy matching | X |  |  |  |
| Aligns names | X | X | X | X |
| Updates names | X | X | X |  |
| Updates synonyms | X | X | X | X |
| Handles phrase names | X |  |  |  |
| Identifies taxon rank of name | X | X | X |  |
| Handles infraspecific taxa | X | X | X | X |
| Handles higher taxonomy | X | X | X | X |
| Handles taxonomic splits | X |  |  |  |
| Retains field note tags | X |  |  |  |
| Available via R package | X | X | X | X |
| Available via browser | X |  | X | X |

**Table 2.**
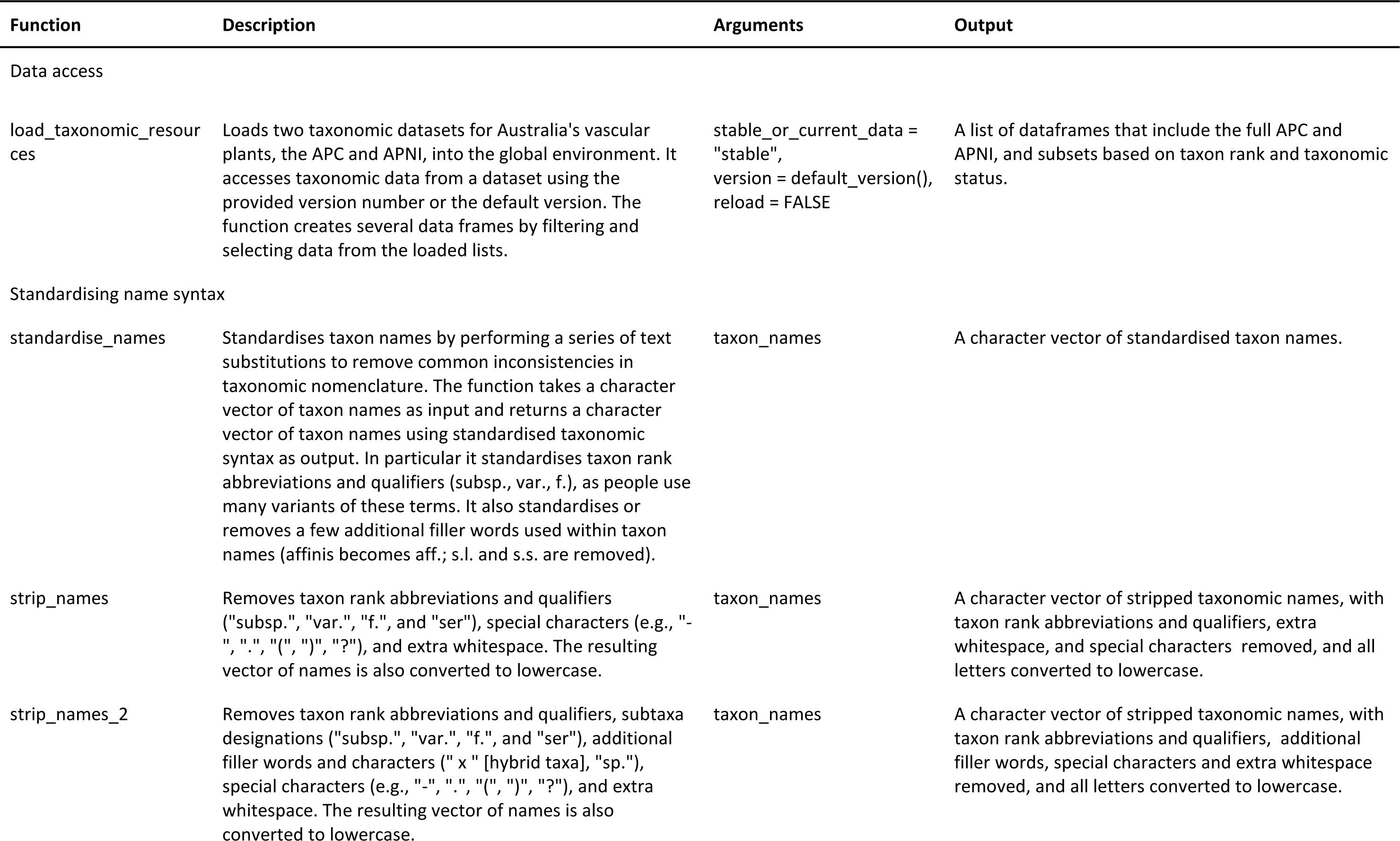
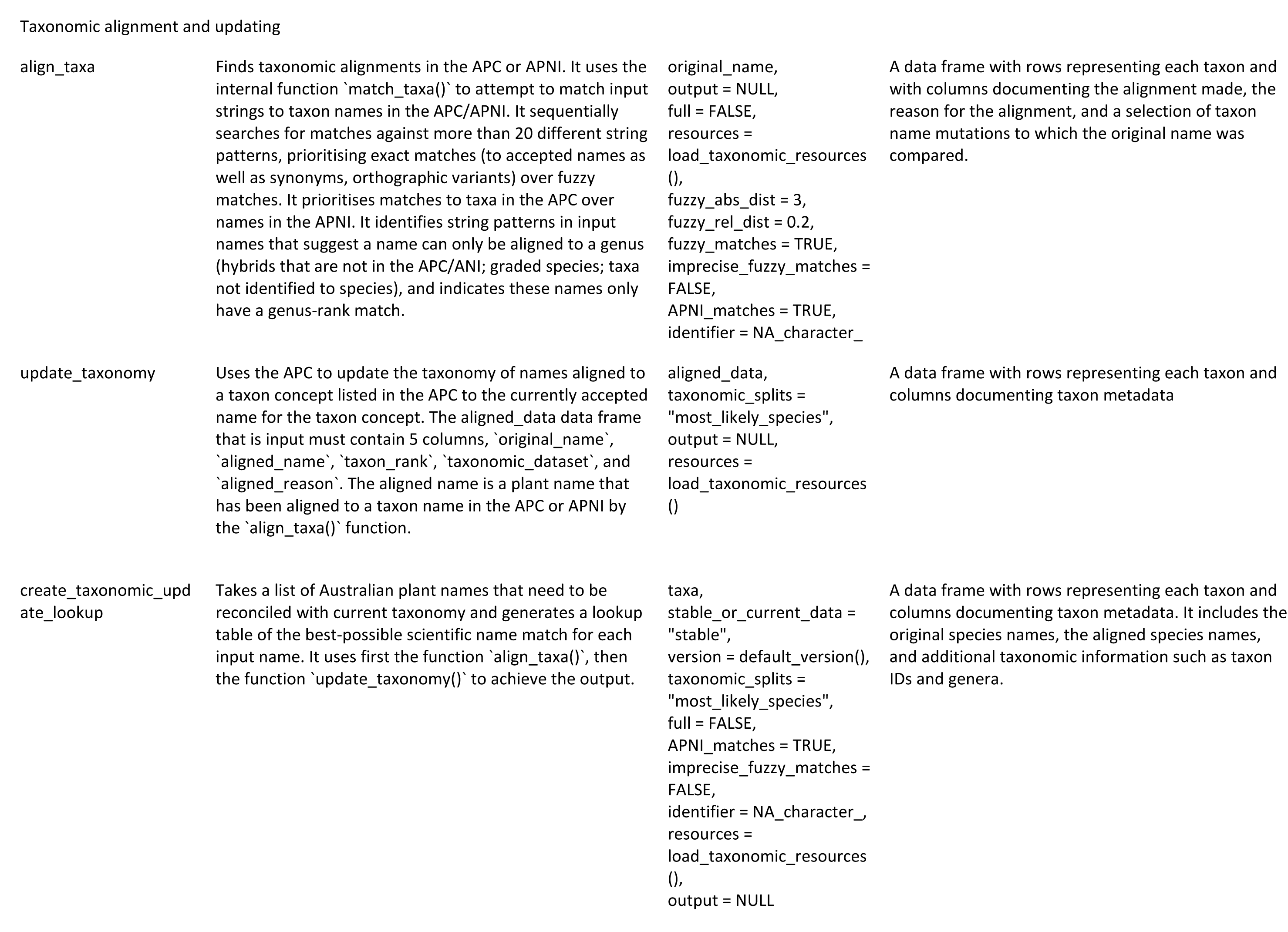
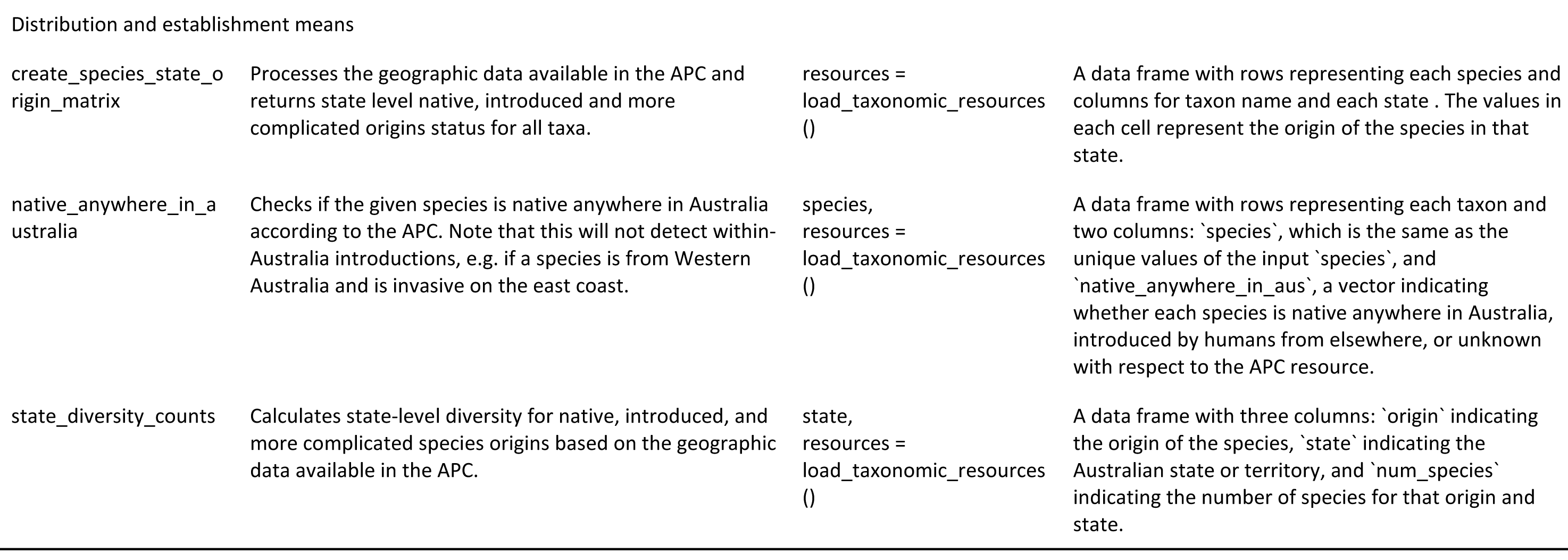
Functions exported by the “APCalign” R package. For additional information see https://traitecoevo.github.io/APCalign/articles/function_notes.html.

### Load taxonomic resources

The taxonomic resources underpinning “APCalign” are the datasets within Australia’s National Species List (auNSL) pertaining to terrestrial vascular plants, the Australian Plant Census (APC) and the Australian Plant Names Index (APNI). The APC is a compilation of endorsed taxon concepts, including both currently accepted taxon concepts and taxon concepts with an alternative taxonomic status, such as synonyms or orthographic variants. Nearly all taxon concepts with a taxonomic status other than “accepted” can be matched to an accepted taxon concept through an identifier (acceptedNameUsageID). For “APCalign”, taxon concepts (and their associated scientific names) within the APC are divided into accepted taxon concepts and taxon concept with an alternative taxonomic status, such as taxonomic synonym, basionym, orthographic variant, and misapplied .Taxon concepts within the APC that have a taxonomic status other than “accepted” are collectively referred to hereafter as ‘APC-synonym’. The APNI is a dataset of all scientific names documented as having been applied to a native or naturalised plant growing in Australia, including both the ∼110,000 names included in the APC and ∼24,500 names not linked to an APC taxon concept. For “APCalign”, only the subset of APNI names not also in APC is considered.

The function ‘load_taxonomic_resources()’ loads copies of the APC and APNI, either a default version (the latest GitHub accessioned version), the latest version, or a specified version number. The function creates several data frames by filtering and selecting data from the loaded lists, and dividing the resources based on the taxon concept’s taxonomic status (for APC) and the taxon rank of the taxon concepts (for APC) or scientific name (for APNI).

### Taxonomic alignment and updating

#### Overview

Aligning the original scientific names to the best match within the APC/APNI requires multiple considerations, including 1) specific syntax patterns that identify the rank to which the name can most appropriately be aligned; 2) whether matches are best made to the entire string or a subset of words; 3) when matches should be made to names from the APC versus APNI; and 4) whether fuzzy matches are appropriate. Users can select whether 1) matches are sought in both the APC and APNI (default) or just the APC; 2) whether fuzzy matching should be enabled (default) or disabled; 3) the initial fuzzy-matching threshold; and 4) whether a second round of imprecise fuzzy matching should be enabled or disabled (default).

Scientific names matched to a taxon concept within the APC are then updated to a currently accepted taxon concept within the APC and additional metadata columns propagated depending on the scientific name’s taxon rank and the taxonomic dataset to which an alignment was made. Users can select how to update names for taxon concepts where taxonomic splits introduce ambiguity in which currently accepted taxon concept is correct.

“APCalign” includes three functions to standardise taxon names (standardise_names()’, ‘strip_names()’ and ‘strip_names_2()), one function to align input names with names in the APC/APNI (align_names()), and one function to update names to those for currently accepted taxon concepts (update_taxonomy()) (Table 2). However, most users will instead use the function ‘create_taxonomic_update_lookup()’ which uses these individual functions to standardise, align, and update the original names in a single step (Tables 2, 3). Example inputs, outputs and code are available at https://traitecoevo.github.io/APCalign/articles/APCalign.html and https://traitecoevo.github.io/APCalign/articles/function_notes.html.

**Table 3.**
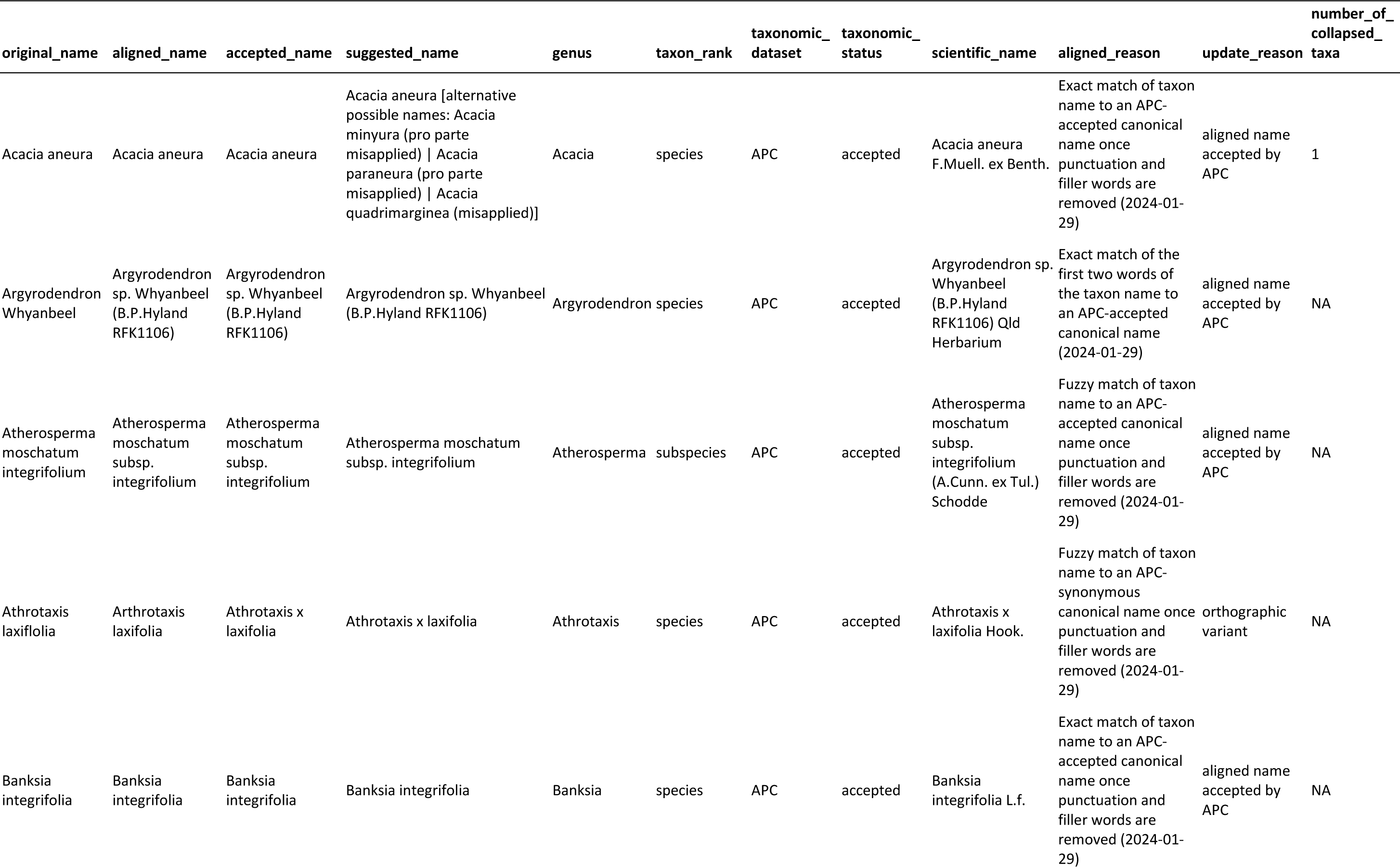
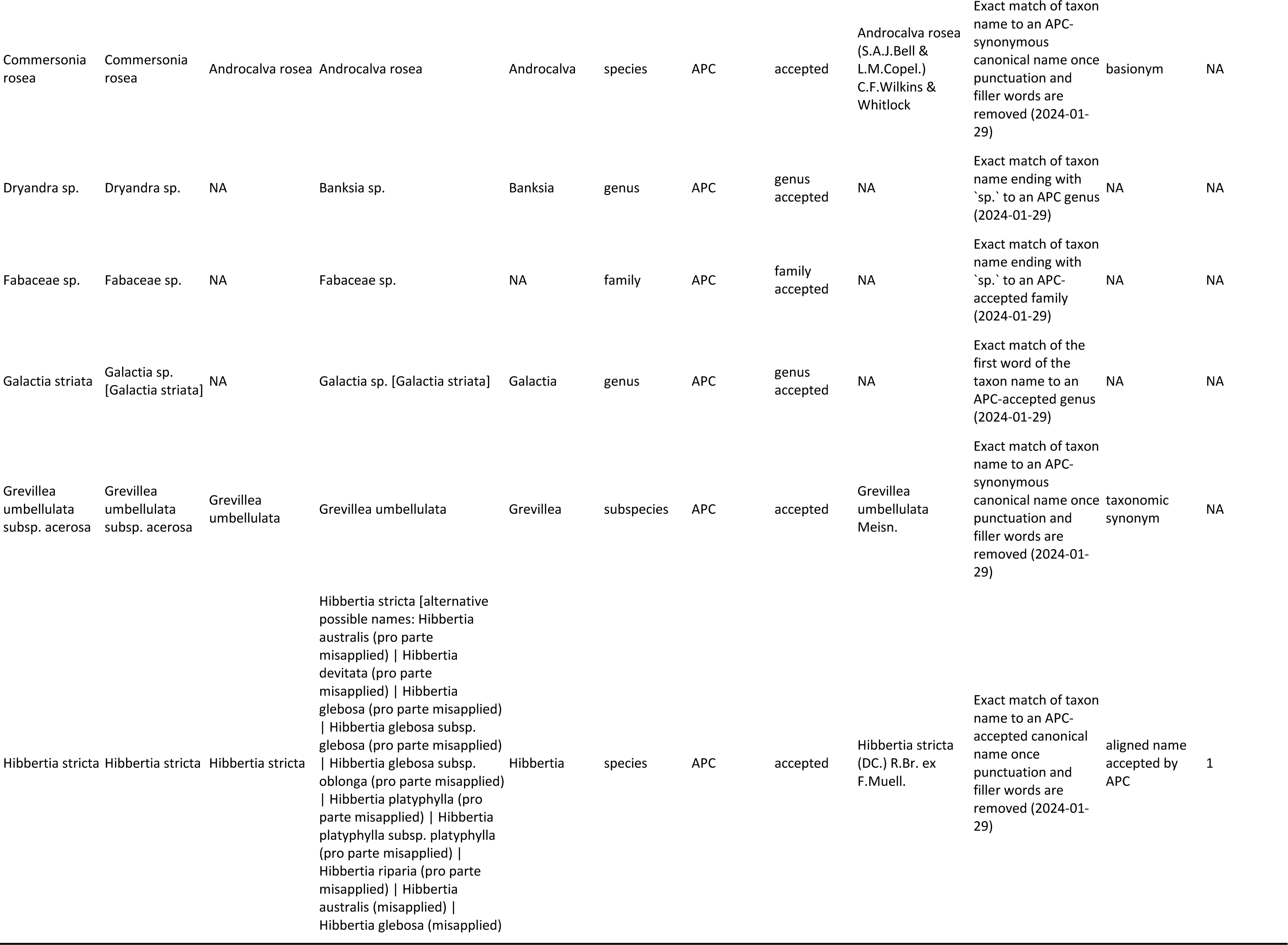
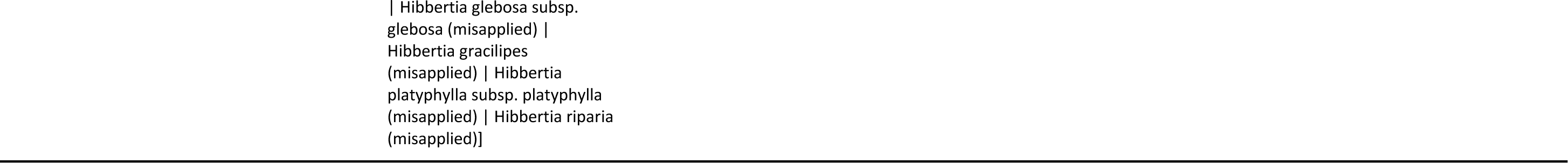
Illustration of outputs from the function ‘create_taxonomic_update_lookup()’ for a series of nomenclatural challenges; the argument ‘taxonomic_splits’ is set to "most_likely_species". For details on *Hibbertia stricta* see (Toelken and Miller 2012).

#### Standardising names

A preprocessing step before aligning input names to names in the APC/APNI is to create six derivations of the original name (for the list of submitted names) or the canonical name (from the taxonomic resources), as different algorithms seek matches against different derivations. For instance, for the APC, matches are sought to scientific names with authorship, scientific names without authorship (termed canonical names), and subsets of words from the canonical name. To aid with name alignments, three standardisation and manipulation functions are applied both to the taxonomic resources and the submitted original names. First, for the original names, taxon rank abbreviations and qualifiers are standardised (to: sp., var., subsp., f., aff., ser.), and variants on *sensu lato*, *sensu stricto*, and excess white space are removed; this is performed by the function ‘standardise_names()’ (Table 2). The output, termed the ‘cleaned_namè, is then passed through one of two additional functions: ‘strip_names()’, which removes punctuation, var., subsp., f., and ser., or ‘strip_names_2()’, which removes both the same characters as ‘strip_names’ and also the filler words cf., sp., and x) (Table 2). From the output of ‘strip_names_2’ one can then extract the first 1, 2 or 3 words, and be confident they are informative terms pertaining to the taxon name. All canonical names in the APC and APNI are also passed through ‘strip_names()’ and ‘strip_names_2()’ to create derivations of the complete scientific names to permit comparisons between the input names and the names in the taxonomic resources.

#### Aligning with APC/APNI names

The function ‘align_taxa()’ then uses the six name derivations to indicate: 1) the lowest taxon rank to which the original name could be aligned (family, genus, species, or infraspecies); 2) the taxonomic dataset to which the original name could be aligned (APC or APNI); 3) for names aligned to the APC, whether the taxonomic status of the aligned name is accepted or a “synonym” (broadly defined as a true synonym or a basionym, orthographic variant, etc.); and 4) the best possible match to a taxon concept (in APC) or other scientific name (in APNI) (Table 4).

**Table 4.**
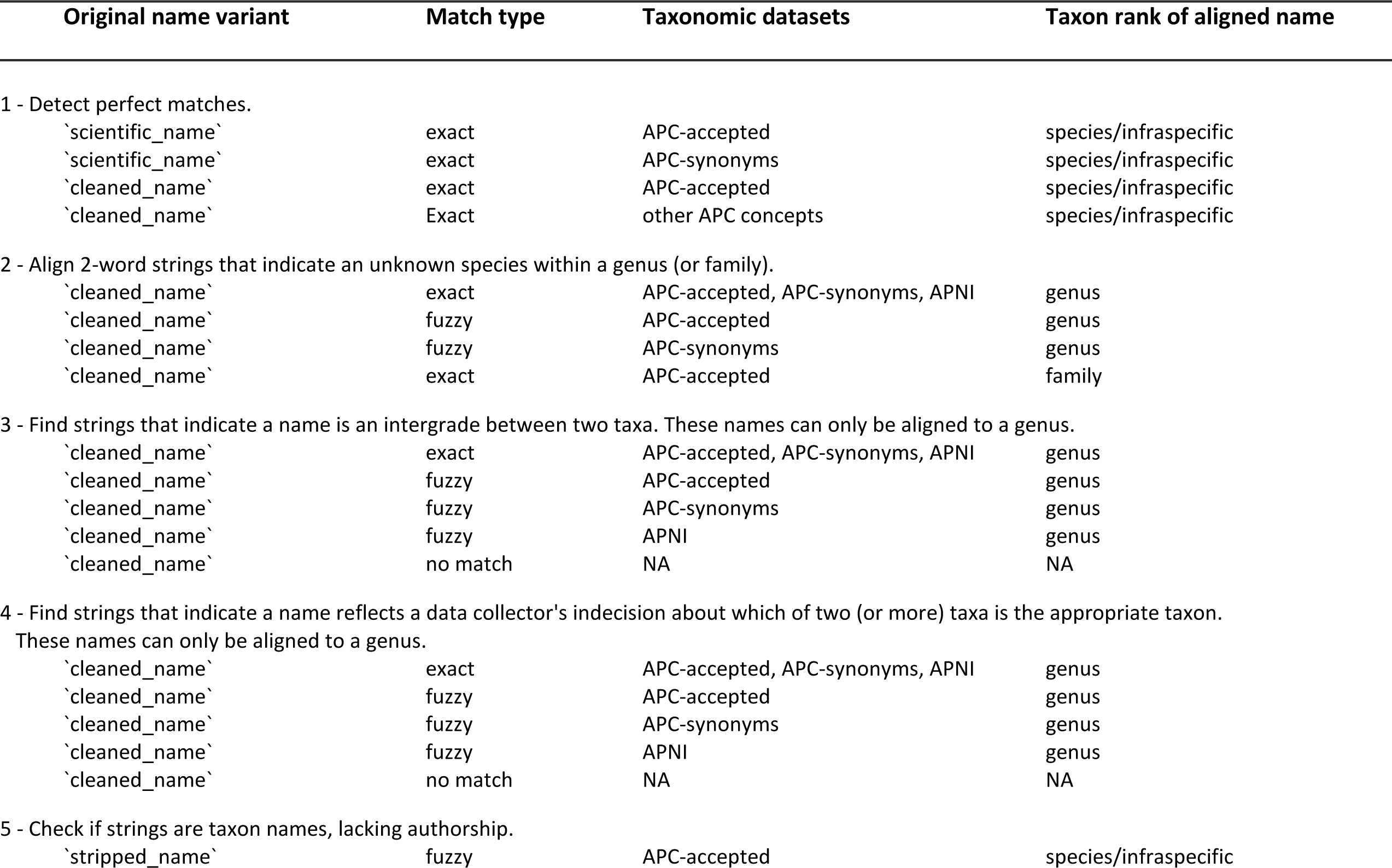
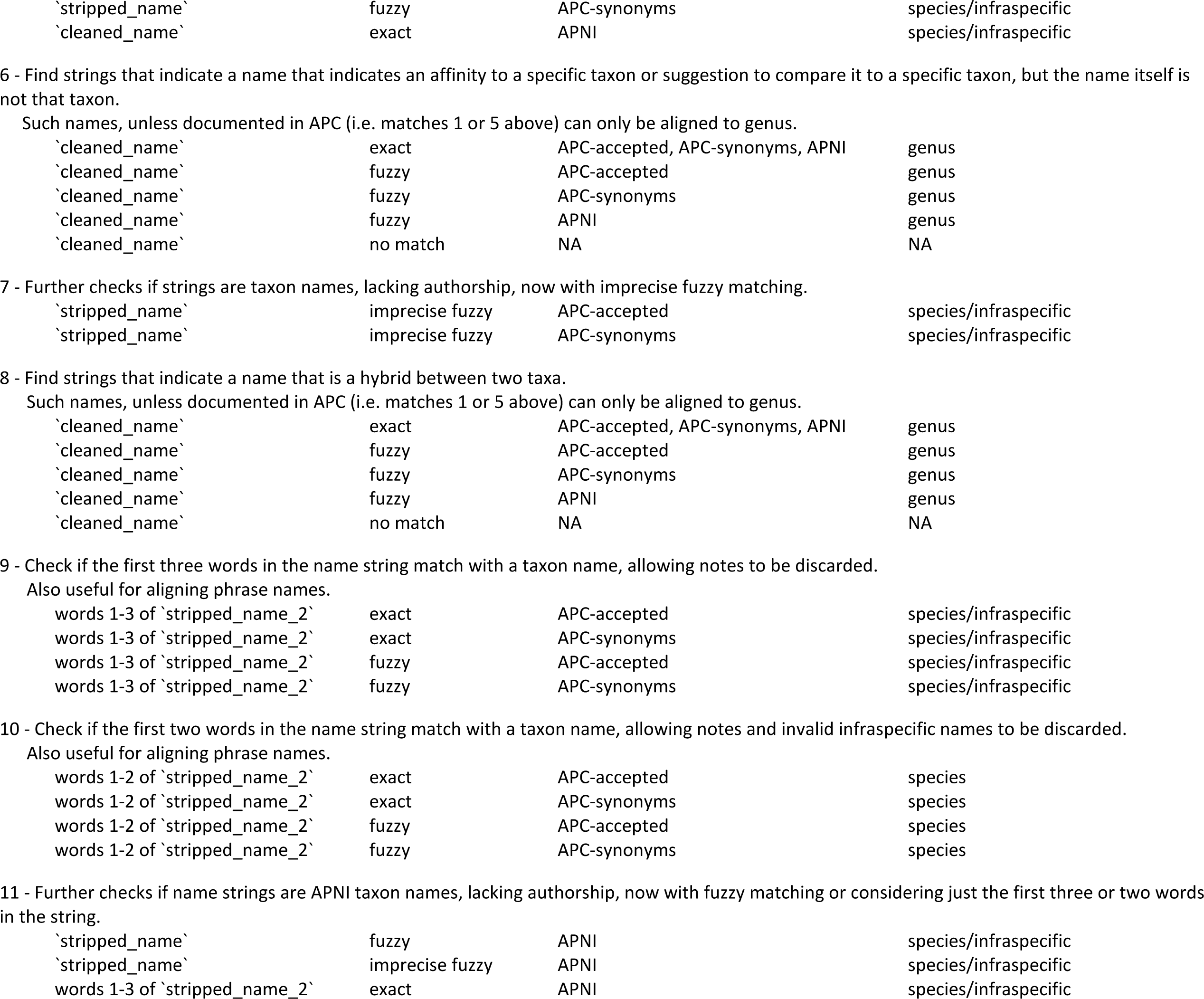
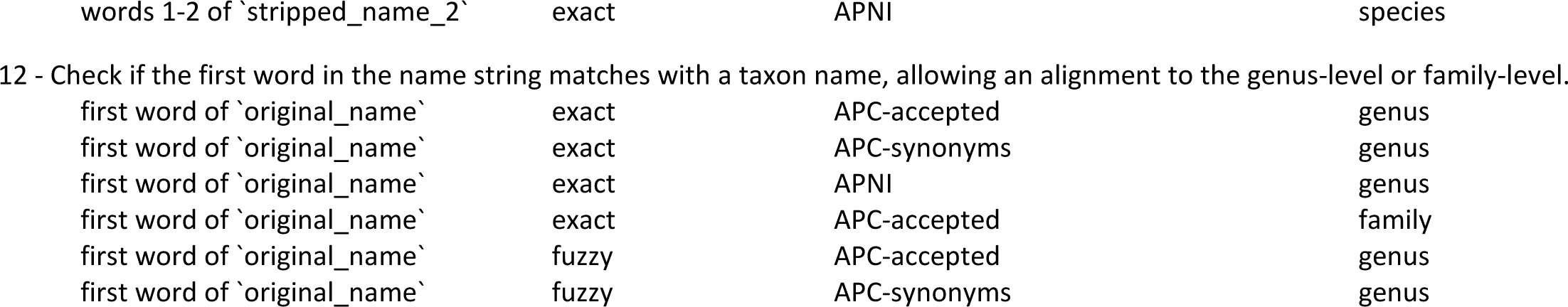
Sequences of queries used in the algorithm to align taxa with known names in APC and APNI. Names are sequentially tested until successful alignment is achieved. The APC is divided into two subsets, one including all accepted taxon concepts (‘APC-accepted’) and a second with taxon concepts with a taxonomic status other than “accepted” (‘APC-synonyms’). For additional information see https://traitecoevo.github.io/APCalign/articles/updating-taxon-names.html.

Perfect matches to scientific names in the APC (with or without authorship) are sought first. If a perfect match is not detected, each original name, or derivation thereof, is automatically, sequentially passed through as many of the additional 47 algorithms as is required until an alignment can be made (Tables 3, 4). Each of these algorithms uses a unique combination of 1) names from the APC/APNI, 2) entire versus partial strings (1, 2 or 3 words), 3) perfect versus fuzzy matches and 4) specific fuzzy match settings. In addition, specific characters are identified that reliably indicate when an original name can never be resolved more specifically than a genus-level (or even family-level) name. The sequence of these match-steps was calibrated to ensure that perfect matches take precedence over fuzzy matches, APC matches over APNI matches, and that taxon names that cannot be aligned to a species are only matched against a list of genus and family names. The sequence of algorithms was carefully curated to maximise the number of accurate alignments by repeatedly testing the 47,000 unique original names from the AusTraits plant trait database (Falster *et al.* 2021). Particular considerations incorporated into the algorithms include:

#### Exact matches of the entire string to the APC and APNI

The first four algorithms detect perfect matches to scientific names in the APC (with or without authorship) (Table 4). Exact matches to APNI names are performed only much later in the series of algorithm steps, as APNI includes many scientific names that are slight spelling variants of the names of taxon concepts documented by the APC, but which have not yet been matched to an APC taxon concept. An explicit decision was made to match these names to a similar name in the APC instead of an exact match in APNI, as otherwise many apparent typographic errors were aligned with a name only present in APNI, rather than an accepted taxon concept in the APC that differed by a single character.

#### Genus-resolution names

Three string patterns reliably indicate whether a name can only be aligned to a genus: 1) “genus sp.” Indicating a plant has only been identified to genus level; 2) “—” indicating an intergrade, and 3) “/” indicating indecision between two names (Table 4). If these string patterns are detected, the name is reformatted as “genus sp. [originally submitted name]”. The function ‘align_taxa()’ includes an argument ‘identifier’ that allows a string to be added to each reformatted name: “genus sp. [originally submitted name; identifier]”. This ensures that, for instance, multiple records of “Eucalyptus sp.” that are documented at different locations or by different people, have an identifier linked that clarifies they are not necessarily identical taxon concepts, but rather members of the genus *Eucalyptus* that the dataset collector could not identify to species level.

There are additional string patterns that can occur either as part of a name documented by the APC/APNI or which indicate a name that can only be resolved to genus level. These include hybrids (detected by the presence of “ x ”), *species affinis* (detected by variants of “aff”), *conferret* (meaning “compare to”, detected by “cf”). If an input name includes these strings, a match is first sought against the APC/APNI; if that fails, genus-level matches are sought. Overall, for all names for which a genus-resolution match is appropriate, the first word of the name is matched sequentially against APC-accepted genera, then ‘APC-synonymous’ genera and APNI genera, followed by fuzzy matches to genera from each of these lists, and finally to APC-accepted family names.

#### Fuzzy matches

Minor typographic errors are commonplace in scientific names in ecological datasets, most often due to incorrect vowel combinations or omitting/incorrectly including double consonants. Fuzzy matching allows such errors to be corrected, but fuzzy match algorithms must be carefully calibrated to correct mistakes without making erroneous matches. There are, therefore, two levels of fuzzy matches performed by the algorithm. For the first, the default is set to a conservative maximum change threshold of 3 characters and 20% of characters, which is sufficient to catch most minor typographic errors. The first letter of each word in the genus, species epithet and, when relevant, infraspecific epithet, remain fixed, as erroneous changes to the first letter of the species epithet are uncommon in datasets, yet allowing the first letter to be changed repeatedly introduced erroneous matches in test datasets. Arguments in the function ‘align_taxa()’ allow the degree of fuzziness to be changed or to disable all fuzzy matches. Later in the algorithm sequence, alignments are sought through less precise fuzzy matches, which allow changes of up to 5 characters or 25% of the characters. This occasionally corrects legitimate typographic errors, but more frequently makes erroneous alignments and is therefore turned off as a default. Each of these function arguments can be adjusted by users, or, if omitted, the defaults are applied.

#### Matches to partial strings

Additional matches (exact or fuzzy) can be accomplished by extracting just particular words from the original name string. Such matches are made on name strings that have been processed by the function ‘strip_names_2()’. Three categories of names are readily aligned when just 2 or 3 words are considered: phrase names, accepted species names with infraspecific epithets that do not exist in the APC/APNI, and names where the “name” includes both a valid scientific name and field notes. Phrase names are accurately aligned through these algorithms, as the first word (usually the genus) and the second and third words (usually a place name or descriptive term; or a place name or descriptive term + voucher number) are generally accurately recorded in the original name, but the exact syntax and the inclusion/omission of the collectors name and initials rarely follows the exact APC convention (Barker 2005). For instance, these matches successfully align “Argyrodendron Whyanbeel” to “Argyrodendron sp. Whyanbeel (B.P.Hyland RFK1106)”, the latter an APC-accepted phrase name (Table 3).

#### Genus-resolution names, part 2

Finally, at the end of the match algorithm sequence, any taxon names that have not been aligned are assumed to only be able to be aligned to genus level. For these, the first word in the original name is matched to lists of genus and family-level APC/APNI names.

The output from the ‘align_taxa()’ function is a table that includes an aligned name, the taxonomic dataset for the aligned name, and the taxon rank of the aligned name.

#### Updating taxonomic names

The function ‘update_taxonomy()’ performs a series of manipulations, including 1) updating names that are “APC-synonyms” to their currently accepted taxon concept (accepted_namè), 2) offering a ‘suggested_namè for each ‘aligned_namè (especially important for ‘aligned_names’ for which there is no ‘accepted namè, as occurs if the ‘aligned_namè is genus-rank or aligned to an APNI name), 3) adding taxon name identifiers to each name, and 4) offering suggested alternative names for taxon names that are ambiguous due to taxon concept splits. The function first splits the table of aligned names into subtables based on the taxonomic dataset and taxon rank specified. This permits different subfunctions to be used to update the taxon names in each subtable, as different rules for updating names and adding taxon concept/scientific name metadata are applied to each group (Table 5).

**Table 5.** Functions used to update aligned scientific names to the currently accepted scientific name, depending on the taxonomic dataset and taxon rank of the aligned name. For additional information see https://traitecoevo.github.io/APCalign/articles/updating-taxon-names.html.

| Function name | Categories of aligned names processed |  | Updates to `aligned_name` | Format of `suggested_name` | Columns filled in |  |  |
| --- | --- | --- | --- | --- | --- | --- | --- |
|  | taxonomic dataset | taxon rank |  |  | accepted name** | scientific name** | genus** |
| update_taxonomy_APC_genus | APC | genus | to APC-accepted genus | genus sp. [notes] | no | no | yes |
| update_taxonomy_APNI_genus | APNI | genus | none | genus sp. [notes] | no | no | no |
| update_taxonomy_APC_family | APC | family | none | family sp. [notes] | no | no | no |
| update_taxonomy_APC_species_and _infraspecific_taxa | APC | species* |  | APC-accepted species name | yes | yes | yes |
| taxonomic_splits = "most_likely_species" |  |  | to APC-accepted taxon concept | most likely APC-accepted species name [alternative possible names] | yes | yes | yes |
| taxonomic_splits = "return_all" |  |  | to APC-accepted taxon concept | all possible APC-accepted species name (extra rows added) | yes | yes | yes |
| taxonomic_splits = "collapse_to_higher_taxon" |  |  | collapsed to APC-accepted genus | genus sp. [collapsed names] | no | no | yes |
| update_taxonomy_APNI_species_and _infraspecific_taxa | APNI | species* | none to `aligned_name`; genus to APC-accepted genus if possible | APNI listed species name | no | yes | sometimes |
| (not aligned) | (not aligned) | (not aligned) | none | none | no | no | no |

\* includes species and infraspecific ranked names
\*\* includes taxon identifiers associated with accepted taxon concept, scientific name, or genus name

The accepted name column is only filled in for species (and infraspecific taxon concepts) that occur in the APC. For taxon concepts whose taxonomic status is “accepted”, their ‘accepted_namè is identical to their ‘aligned_namè. Names that have been aligned to the list of ‘APC-synonyms’ are linked to taxon concepts with a taxonomic status other than “accepted”. The APC includes a column ‘accepted_name_usage_ID’ that allows ‘APC-synonym’ names to be matched to the currently accepted taxon concept; this column makes it possible to update ‘APC- synonym’ aligned names to the appropriate APC-accepted name (and corresponding taxon concept).

The ‘suggested_namè column offers a best-practise name for all input names that can be aligned at least at the family level. In order of preference, the suggested name is an APC-accepted taxon concept, a taxon concept from the ‘APC-synonym’ list which has not been connected to an accepted taxon concept in the APC (a rare situation), a documented APNI canonical name (the aligned name), or a genus-resolution or family-resolution aligned name. For genus-resolution names, where the genus is a ‘synonym’ (broadly defined) of an ‘APC-accepted’ genus, the genus is updated to the currently accepted genus name.

When possible, unique identifiers provided as part of the auNSL resources are attached to each taxon concept (for species and infraspecific taxon concepts aligned to the APC), scientific names (for all species and infraspecific names aligned to either the APC or APNI), and the genus taxon concept (where the aligned genus is accepted by the APC).

For APC-accepted taxon concepts, taxonomic splits offer an additional complication. A taxonomic split occurs when a previously unified taxon concept has been divided into multiple unique taxon concepts. When the aligned name is the name of the older taxon concept, it is ambiguous whether this name refers to the older broader taxon concept or the current, more narrowly circumscribed taxon concept. The ‘update_taxonomy()’ function includes an argument ‘taxonomic_splits’ that offers three different outputs for these scenarios: 1) the default is to return the name of the “most_likely” taxon concept, the taxon concept that existed in the past as a broad taxon concept and continues to be used today as a narrower taxon concept. The list of alternative taxon concept names is included in square brackets following the most likely name (Table 3); 2) “collapse_to_higher_taxon” aligns such names to the genus, implying that is the best resolution that can be achieved for these ambiguous taxon concepts. The list of currently accepted taxon concept names is included in square brackets following the genus; 3) “return_all” leads to a longer output table with all currently accepted taxon concepts returned as separate rows.

### Distribution and establishment means

The APC also specifies state-level distribution for the accepted taxa, including information about their native or introduced status; this piece of data is termed “establishment means” in DarwinCore (Wieczorek *et al.* 2012). Information about state-level establishment means is a result of a complex consultation process involving the state herbaria. A static version of this is available (Martín-Forés *et al.* 2023) along with R scripts. A very similar product is available here, but in a more user-friendly R package with automatic file handling that also dynamically updates as APC moves forward.

One complex feature of the Australian flora is internal out-of-range introductions, with taxa from one part of the continent becoming naturalised in another (Figure 3). Some of these taxa become very successful outside of their “native” range. A specific type of introduction that may become more important under future climate scenarios is where taxa from mainland Australia have been very recently introduced to Tasmania and other islands. The full complexity of taxon distribution and native status documented in the APC data is available via two functions ‘create_species_state_origin_matrix()’ and ‘native_anywhere_in_australia()’. The function ‘create_species_state_origin_matrix()’ can be especially informative for specific taxa (Figure 3). For the reduced species lookup table, treating all of Australia as an entity, there is a function ‘native_anywhere_in_australia()’. A related function, ‘state_diversity_counts()’ calculates state-level diversity for native taxa, introduced taxa, and taxa with more complicated origins (Table 2).

### Performance and comparison with existing tools

The “APCalign” algorithms were explicitly developed to standardise and align diverse lists of taxonomic names submitted to the AusTraits plant trait database (Falster *et al.* 2021). These taxon lists were derived from ecological field studies, taxonomic revisions, online floras, experimental datasets, and government biodiversity reports. Each category of dataset offered unique challenges that were considered in algorithm development. In particular, aligning the many phrase names in common use in Australian taxonomic literature required distinct tools not yet available in any other existing taxonomic resolution tools worldwide. Although the APC prescribes an exact syntax for phrase names, such as the absence of spaces between the collector’s initials (Barker 2005), phrase names in different resources rarely align perfectly with the APC and fuzzy matches cannot reliably match names with missing collectors or voucher numbers. The solution within “APCalign” was, late in the algorithm sequence, to perform matches on just the first two or three “real” words.

To test the performance of “APCalign” across diverse datasets, the 45,337 unique taxon names contained within the 363 datasets in AusTraits v4.2.0 were submitted to “APCalign”. The distinct names in the AusTraits dataset include minor syntax variants within the dataset (e.g. ‘subsp.’ versus ‘ssp.’), more significant syntax variants (absence of taxon designations such as ‘subsp.’ and ‘var.’; incomplete phrase names), spelling mistakes (in the original datasets submitted to AusTraits), taxonomic synonyms, and incomplete names (e.g. Acacia sp. [long leaves]). 95.7% of the AusTraits names had perfect matches to names in the APC (whether accepted or synonyms, with or without authorship) (Figure 1); the effort to create alignment algorithms focused on the final 4.3% of names.

Most of the additional matches were names where a conservative fuzzy match (changes of fewer than 3 characters and 20% of characters) aligned the name to the APC (1.8%) or names that were documented in the APNI (0.4%). However, the algorithms for other alignments captured important classes of names. In particular, many ecological datasets include names that indicate a taxon that the observer could not resolve to species, with names that follow some variant of ‘genus sp.’ or ‘genus species A/species B’ or ‘genus sp. (notes)’.

A sequence of different algorithms was used to identify all genus- and family-level names and confirm the genus was listed within the APC or APNI. 0.7% of all taxon names could only be resolved to genus- or family-level names. The matches that looked at only the first two or three words, ignoring all infraspecific taxon designations and punctuation, also proved essential to accurately align many taxon names documented in AusTraits datasets. These matches aligned an additional 0.8% of taxa to an accepted name, mostly aligning 1) two or three words from the submitted name string where the name string included notes and 2) phrase names. Just 255 names (0.1%) could not be aligned by any of the matching algorithms. Importantly, of the 99.9% of names that were aligned, there were <0.1% that the AusTraits team deemed to be incorrect alignments during manual checks; these were almost entirely “imprecise fuzzy matches”, which are known to often introduce errors and are, by default, turned off within the “APCalign” functions. When turned on, 0.1% of taxon names were aligned by the imprecise fuzzy matches but about half of these were subsequently reverted, indicating they are a useful tool for exploring possible name alignments, but not for mass processing when each aligned name will not be reviewed.

We also compared the alignment algorithms of “APCalign” with those of three existing global tools: “TNRS” (Taxonomic Name Resolution Service; Boyle *et al.* 2013), “taxize” (Chamberlain and Szöcs 2013; Chamberlain *et al.* 2022), and “kewr” (Walker 2021). “taxize” describes itself as a taxonomic toolbelt, simultaneously searching 20 separate global taxonomic resources to find matches, while “kewr” matches names to 6 different taxonomic resources. There is a significant overlap between these resources. “TNRS” uses the World Flora Online (Borsch *et al.* 2020) and the World Checklist of Vascular Plants (Govaerts *et al.* 2021). These R-based tools all allow lists of names to be read in, offering the rapid alignment of entire datasets. Across the Tree of Life, many such packages exist, referencing either global or regional taxonomic datasets (Boyle *et al.* 2013; Grenié *et al.* 2023).

Of the tools compared, “APCalign”, “TNRS”, and “taxize” include fuzzy matching algorithms, required to correct the minor typographic errors that proliferate in scientific names (Table 1). Meanwhile, “kewr” does not perform syntax standardisation or fuzzy matches and therefore returns ‘NA’ when an exact match is not detected. Another feature of “APCalign” that is shared by “TNRS” and “taxize” is an output column that identifies the resolution of the output name, indicating, if, for instance, a name can be matched to a genus or family, but not a species (Table 1). The outputs from each tool vary according to the encoded matching algorithms. While “TNRS” and “taxize” also incorporated fuzzy matching, a distinctive feature of the “APCalign” fuzzy matching function is that the first letter of each word cannot change. This occasionally misses a valid alignment, but without this feature there are many additional false alignments made. “APCalign” also allows the fuzzy-matching level to be adjusted, should a user wish to institute more or less conservative fuzzy-matching.

Another feature of “APCalign”, not shared by the other tools, was detecting a series of symbols/punctuation that indicated the name could only be aligned to a genus, not a species. These include “x” (a hybrid), “aff.” (given taxon shows an affinity to the named taxon) “cf.” (given taxon should be compared to the name taxon), “--” (taxon is an intergrade between two taxa), and “/” (two taxon names submitted, because the author is unsure which is correct). The comparison tools all prioritise looking at just the first words in the string, which leads to the submitted name being aligned to the first of the names listed, while “APCalign” automatically flags these names as only being appropriate to align to a genus. The initially submitted string is then inserted in square brackets following the genus, so the information is not lost (e.g. ‘genus sp. [genus species x genus species]’). In APC align the 2-word and 3-word matches are only made after names with such syntax patterns are flagged as only being suitable to align to genus-resolution, and hence are not passed through these matching algorithms.

One “APCalign” feature that is not available in any of the other outputs is the addition of a custom identifier for names resolved to the genus- or family-level. This does not improve alignment or increase our taxonomic interpretation of a specific name but does offer a means to indicate that a taxon name that is only resolvable to a particular genus or family represents an unidentified species within a given dataset or location. For AusTraits, this is essential, as many datasets have submitted the name ‘Acacia sp.’ and a mechanism is required to indicate these do not all refer to the same taxon concept, but instead to a multitude of different *Acacia* species that the dataset contributor could not identify to species.

An important feature of “APCalign” shared by “TNRS” and “taxize” is the ability to update outdated taxon concepts to the names associated with their currently accepted . As lists of taxonomic names quickly become outdated it is time-consuming to manually determine which names must be updated, then change them one-by-one. As taxonomic concepts are a moving target, different taxon lists will consider different names to be “current”. As such, while “TNRS” and “APCalign” agree on the currently correct name for the majority of taxa, there are instances where “TNRS” returned a recently updated name, while “APCalign” reported an older name that is still considered current by the APC. This asynchrony will always exist across taxonomic lists, as each is updated by a different panel who, for some taxa, will make divergent judgements on the application of each name and review name changes at different times. All detected names that were updated by “TNRS”, but not “APCalign” were manually checked and all were names that had been changed in the past 5 years.

The goal of “APCalign” is to align submitted names to the currently accepted name within Australia’s two vascular plant taxonomic resources, APC and APNI. “TNRS” offers a similar aim, but at a global scale, and is therefore aligned to a global taxonomic resource. In contrast, “taxize” and “kewr”, while seeking to standardise names, have the objective of matching an input taxon name to associated taxon names across many resources. For instance, while the “APCalign” and “TNRS” workflows aim to replace all outdated taxonomy with a currently accepted name, “taxize” and “kewr”’s outputs are developed to display “here are all the names, in all these resources, that are linked to the string that was input”, both leaving more name selection decisions up to the user and providing a snapshot of the complexity of selecting a “best” taxonomic name across all global resources.

### Options for different users

While “APCalign” was created with researchers who are well-versed in R programming in mind, we also wanted to cater to non-R users. For non-R users who want to align and update plant taxonomic names, we recommend using the “APCalign-app” web interface (https://posit-connect-unsw.intersect.org.au/APCalign-app/). “APCalign-app” provides an easy entry point to the core functionality of the R package by replacing the multi-step, code workflow with a simple interface. The interface is preset with intuitive default settings that can be bypassed. This allows users to feasibly supply their taxonomic names, either by typing into an input box or uploading a .csv file and obtain aligned and updated names that can be downloaded. We hope this mobile-enabled web interface can improve the consistency of plant taxonomic names for all users in the botanical community.

## Conclusions

Australian researchers are fortunate to have the resources provided as part of the Australian National Species List for vascular plants, regularly updated to incorporate taxonomic revisions and the names of newly described taxa. This paper presents a new interface to the dynamic APC, which is the consensus taxonomy for native and naturalised Australian plants. It is essential that research projects, biodiversity assessments, and indeed nurseries, use the scientific names for vascular plants accepted by the APC to facilitate communication at the national level.

Country or continental efforts will always be complementary to global efforts. While there exist global taxon-matching tools with similar functionalities, none reference the APC. Our comparison with the global tool “TNRS” highlighted the importance of a local tool, as only the APC includes phrase names and aligns names to those specified within the Australian National Species Lists for vascular plants. Unlike global taxonomic alignment tools, “APCalign” also considers taxonomic splits, allowing the user to decide what scientific name they consider appropriate for taxon concepts that have been subdivided during taxonomic revisions. The Brazil Flora 2020 effort has a similar country-specific tool with a shiny interface (http://www.plantminer.com/), specific to the Brazilian flora. An added benefit of working with a continent- or country-specific list is that fuzzy matching and synonym correction are less prone to false positive errors with smaller lists.

Although “APCalign” was only released as a complete R-package in late 2023, the underlying code has been tested on multiple datasets during its development, including the 47,000+ taxon names submitted to AusTraits, 12600+ taxon names submitted to the AIslands project on the floras of Australia’s islands (J. Schrader, pers. com.), and 15,700+ taxon names within the New South Wales BioNet Atlas (Department of Planning and Environment; https://www.environment.nsw.gov.au/topics/animals-and-plants/biodiversity/nsw-bionet). These projects and others have each merged disparate datasets collected across more than a century, successfully outputting harmonised compilations of updated taxon names. Taxonomy will always be a moving target, but we have built the “APCalign” package to be relatively stable as the APC continues to evolve and improve. We hope that others can use the tool to synchronise their research with the APC, leveraging the actively developed, high quality taxonomic resource that the APC provides.

## Acknowledgements

We thank the ARDC for their commitment to funding projects that increase community access to and use of informatics resources. We thank Rachael Gallagher for many conversations about how to build the best tool for Australia’s research community. We thank David Coleman and Julian Schrader for testing the APCalign functions on two large datasets, offering feedback on needed algorithm refinements.

## Authorship

EW and DF led the development of the algorithms to align and update taxon names. EW, FK, WC, and DF wrote the software. AF, AM, and HS offered expertise to ensure the algorithms correctly aligned and updated all categories of correct and erroneous taxon names. All authors contributed to the writing of the manuscript.

## Funding

The AusTraits project received investment (https://doi.org/10.47 486/DP720) from the Australian Research Data Commons (ARDC). The ARDC is funded by the National Collaborative Research Infrastructure Strategy (NCRIS). F.K. was funded by a UNSW Research Infrastructure Grant to Falster.

## Data Availability Statement

The code underpinning “APCalign” is available on the project’s GitHub repository, https://github.com/traitecoevo/APCalign. Articles on the GitHub repository’s website offer

additional details on the functions, including example usage; see https://traitecoevo.github.io/APCalign. The APC and APNI, the Australia’s National Species List

resources pertaining to vascular plants, can be accessed at https://biodiversity.org.au/nsl/services/export/index.

## Conflict of Interest Statement

The authors declare no conflicts of interest.

## Notes

https://github.com/traitecoevo/APCalign

### Competing Interest Statement

The authors have declared no competing interest.

